# A Large Animal Model of Heritable Pulmonary Arterial Hypertension Using Gene-edited *BMPR2* Sheep

**DOI:** 10.64898/2026.02.06.704456

**Authors:** Sanjeev A. Datar, Nicholas Werry, Austin R. Brown, Devon S. Fitzpatrick, Oluwafemi Falade, Josephine F. Trott, Rachel Hutchings, Elena K. Amin, Jessica M. Morgan, Hythem Nawaytou, Gail H. Deutsch, Eric G. Johnson, Omar A. Gonzales Viera, Thomas F. Bishop, Tara Urbano, Bret R. McNabb, Eric D. Austin, Jeffery R. Fineman, Alison L. Van Eenennaam

**Affiliations:** Department of Pediatrics, University of California, San Francisco, CA 94143, USA; Department of Animal Science, University of California, Davis, CA 95616, USA; School of Veterinary Medicine, University of California, Davis, CA 95616, USA; Department of Laboratory Medicine and Pathology, University of Washington, Seattle, WA 98195; School of Medicine, Vanderbilt University, Nashville, TN 37235, USA

## Abstract

Pulmonary Arterial Hypertension (PAH) is a rare vascular disorder characterized by elevated pressure in pulmonary arteries, eventually leading to right ventricular failure. Approximately 50% of pediatric disease and 20% of adult disease can be linked to a genetic mutation, with nearly 70% of these cases involving mutations in the bone morphogenetic protein receptor type 2 (*BMPR2*) locus. Investigations using rodent models have made significant advances in our understanding of BMPR2 signaling; however, limited data exist regarding the onset and course of PAH, and etiologies for phenotypic expression in these patients remain unknown. In this work, we describe the development of a novel ovine model of heritable PAH. Because homozygous disruption of *BMPR2* is embryonic lethal, we developed heterozygous *BMPR2* sheep by using a PAM-disrupting synonymous single stranded oligodeoxyribonucleotide alongside a single guide RNA and Cas9 mediated gene editing strategy. The resulting *BMPR2^(+/–)^* lambs demonstrated cardiac and pulmonary vascular pathology that are consistent with *BMPR2* mutation-driven PAH observed in humans. Given the genetic and physiological similarities of *BMPR2^(+/–)^*sheep to humans with heritable PAH, this large animal model will serve as a vital platform for mechanistic molecular studies and will provide a much-needed pre-clinical model for extensive treatment evaluations.

## Introduction

Pulmonary hypertension (PH) represents a spectrum of disease processes whose underlying pathophysiology contributes to a common, and often lethal phenotype: elevated pressure in the pulmonary arteries that eventually leads to right heart failure. Detailed investigations have been hampered by the lack of clinically relevant animal models of disease, and in particular, large animal models in which pre-clinical trials can be adequately performed. The 7^th^ WSPH classified pulmonary vascular disorders into five groups and 30 subgroups (1). Group I PH is pulmonary arterial hypertension (PAH). This is the most common diagnosis, with a large subpopulation having heritable disease with a family history of PAH. Over the past decade, the genetic basis underlying many cases of PAH have been identified. Mutations in twelve genes have now been strongly linked to PAH (2); indeed, between 70 and 80% of hereditary PAH and up to 10% of idiopathic PAH can be attributed to mutations in known genes (3). In both children and adults, 70-80% of these cases involve mutations in *BMPR2* (bone morphogenetic protein receptor type 2) (4). *BMPR2* is particularly noteworthy since a missense mutation in exon 3 of this gene is associated with one of the largest familial cohorts of PAH (5, 6). Patients with the *BMPR2* mutation are at high risk for developing PAH, but penetrance is highly variable with pathologic expression occurring in ∼42% of females but only ∼14% of males (7, 8). Its onset can be insidious, despite careful monitoring and follow-up (9), and the natural history of this disease can be quite variable (10).

When functional, *BMPR2* encodes a type II serine/threonine kinase receptor in the transforming growth factor beta superfamily. It forms a transmembrane complex with two type 1 receptors that bind extracellular bone morphogenic protein (BMP) growth factors and induce downstream activation of *SMAD 1/5/8* to regulate survival and propagation of vascular cells (11). Notably, in murine models, complete loss of *Bmpr2* is early embryonic lethal, failing to form organized structures or mesoderm (12), and *BMPR2* loss-of-function mutations have only been found in heterozygous individuals (13).

Sheep have long been regarded as high-fidelity models of newborn and infant physiology (14–20). As CRISPR/Cas9 gene editing has emerged as a routine methodology, genetic manipulation in sheep has now become logistically and economically feasible; indeed, a number of genetic ovine models of human disease have been recently described, including those for human deafness (21) and cystic fibrosis (22). Until now, however, a large animal model of hereditary PAH has not been successfully created, in part due to the challenges of introducing a heterozygous edit thereby allowing the gene-edited fetuses to survive though gestation and parturition.

Many modern approaches to gene editing include the use of single-stranded oligodeoxyribonucleotides (ssODNs). These short DNA sequences serve as customizable donor templates for homology directed repair (HDR), offering high editing efficiency and low incidence of off-target editing (23, 24). By combining CRISPR-Cas9-mediated cleavage and ssODN-mediated repair, it is possible to introduce a desired mutation to a specific site within the DNA

(25). ssODNs have demonstrated remarkable efficiency in driving HDR in zygotes, making them particularly relevant for production of gene-edited animals (26). Importantly, an ssODN can be designed to replace the NGG protospacer-adjacent motif (PAM) with a silent PAM-disrupting sequence that does not alter the protein coded by that sequence, but that prevents all further CRISPR/Cas9 editing at that location (27).

Here, we have coupled a PAM-disrupting synonymous ssODN with a single guide RNA (sgRNA) mediated gene-editing to produce a heterozygous *BMPR2^(+/–)^* ovine model, providing an opportunity to study the molecular pathogenesis of PAH not currently possible in human patients or smaller mammals. The development process of this model represents a substantial accomplishment in livestock gene-editing, while the model itself provides a significant improvement in relevance to human physiology compared to murine studies, and will be an essential research tool for mechanistic understanding, strategies for disease management, and novel therapeutics.

## Results

### HDR Optimization

Using 600 ng/ul ODN, we found that sgRNA1 was significantly more efficient for HDR repair compared to sgRNA2 (p<0.05). The editing efficiency of blastocysts when using the 600 ng/ul ODN and sgRNA1 combination was 44.7% high (>66%), 42.6% intermediate (66-33%), and 8.5% low (<33%) editing, with 4.2% being wildtype. Additionally, the HDR rate was 27.7% high (>20%), 21.3% intermediate (20-5%), and 51% low to no (<5%) HDR. Based on these data, the combination of sgRNA1 with 600 ng/µl ssODN were used to generate *BMPR2^(+/–)^*sheep.

### Gene-Edited Lamb Genotypes

Following electroporation and maturation, five to six blastocysts were surgically transferred to each of eight surrogate ewes that responded to hormone synchronization, and pregnancies were confirmed by ELISA and an ultrasound at 6 weeks post-transfer. A follow-up post-transfer ultrasound at 9.5 weeks confirmed the viability of eight fetuses past the period of embryonic lethality. A total of eight lambs were carried to term. Four lambs were liveborn (3♀ lambs #5, #6, #8; 1♂ lamb #4), and four died perinatally. DNA was collected from all eight lambs and the *BMPR2* region was amplified using Intron2-F2 and Intron-3R primers revealing obvious insertions and deletions in exon 3 of *BMPR2* in three of the eight lambs (Figure 2). The PCR products from each sheep were submitted for both sanger sequencing and Illumina EZ-Amplicon sequencing to determine the alleles present and their relative abundance within each sample (Table 2), revealing at least one wildtype allele generated by HDR in all eight lambs, and at least one edited allele in seven lambs (Table 2; Figure 3). The predicted amino acid sequences of the BMPR2 proteins from edited alleles are in Supplementary Table 2. Seven of the nine edited alleles are expected to result in dramatically truncated BMPR2 proteins, including both the 49 bp and 7 bp deletion alleles of the live heterozygous male lamb #4. All liveborn lambs each carried at least one *BMPR2* knockout allele (Table 2; Supplementary Table 2).

**Figure 1.**
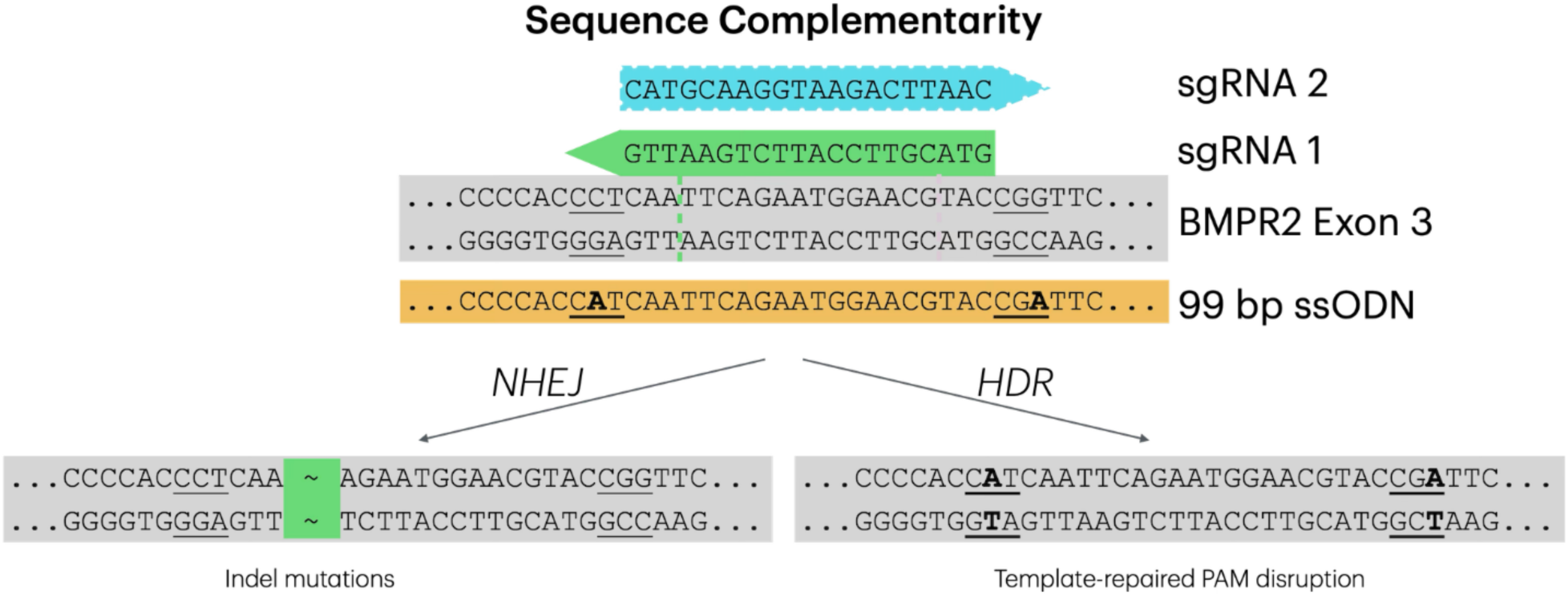
Editing sequence design. Single guide RNA (sgRNA) 1 (green) and 2 (blue) were targeted to exon 3 of *BMPR2.* Protospacer-adjacent motifs (PAMs) are indicated by underlining, mismatches relative to the Oar_v3.1 reference genome are indicated in bold. The location where sgRNA1 would be predicted to introduce a double-stranded break (DSB) is indicated by the green dashed line. Non-homologous end joining (NHEJ) repair of the DSB will repeatedly cleave and introduce indel mutations until the guide sequence is disrupted, while homology directed repair (HDR) using the 99 bp single-stranded oligodeoxyribonucleotide (ssODN) will introduce a synonymous silent mutation disrupting the sgRNA 1 PAM (NGG to NTG) and preventing further Cas9 activity. The additional PAM-disrupting mutation (NGG to NGA) towards the 3’ end of the ssODN, was designed to disrupt the PAM of sgRNA 2.

**Figure 2.**
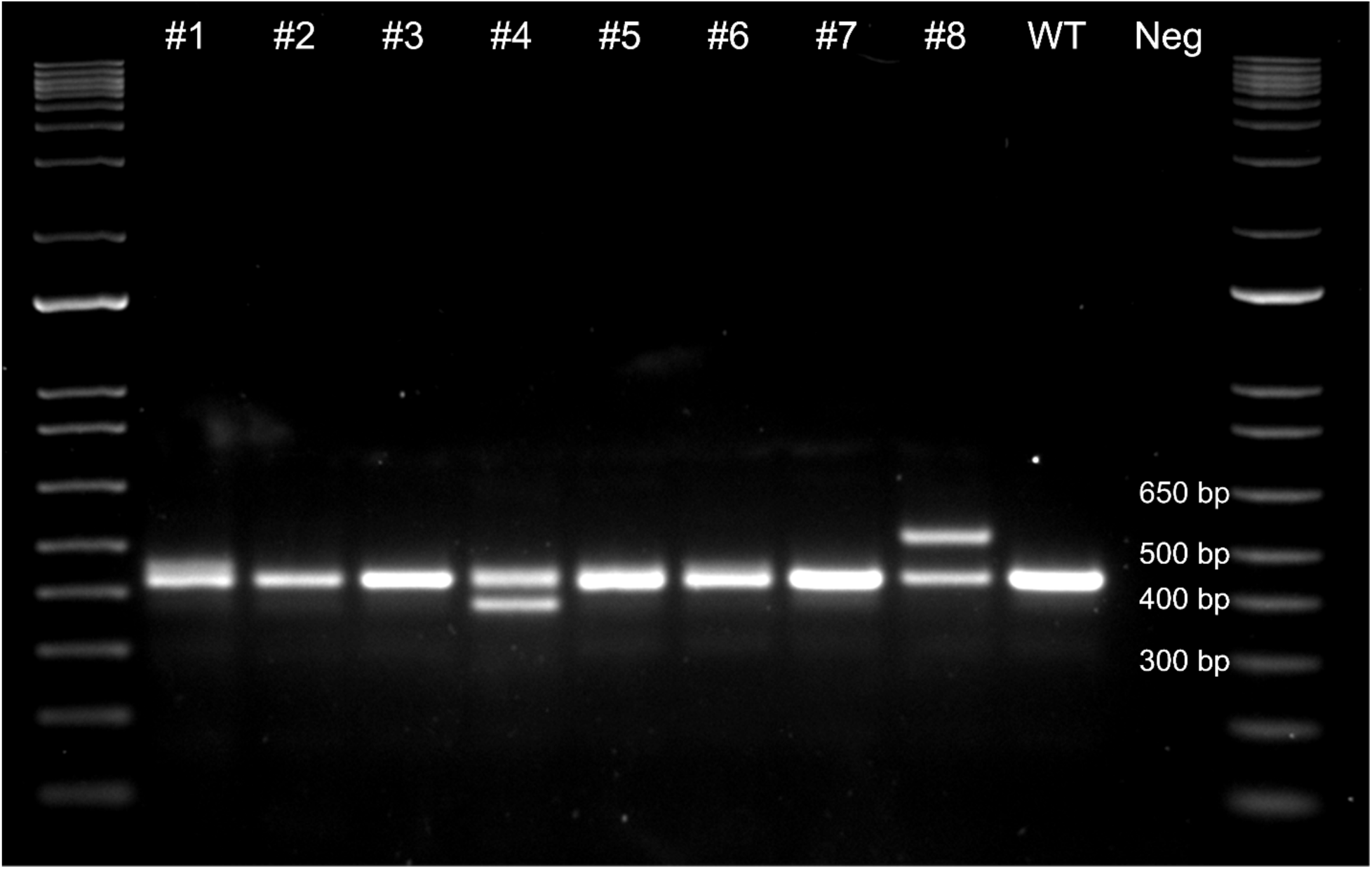
Agarose gel of *BMPR2* lamb genotypes. A 431 bp wildtype (WT) amplicon of *BMPR2* was amplified by primers Intron2-F2 and Intron-3R and visualized on a 1.75% agarose gel to reveal obvious insertions and deletions in several of the 8 lambs. Neg, negative control.

**Figure 3.**
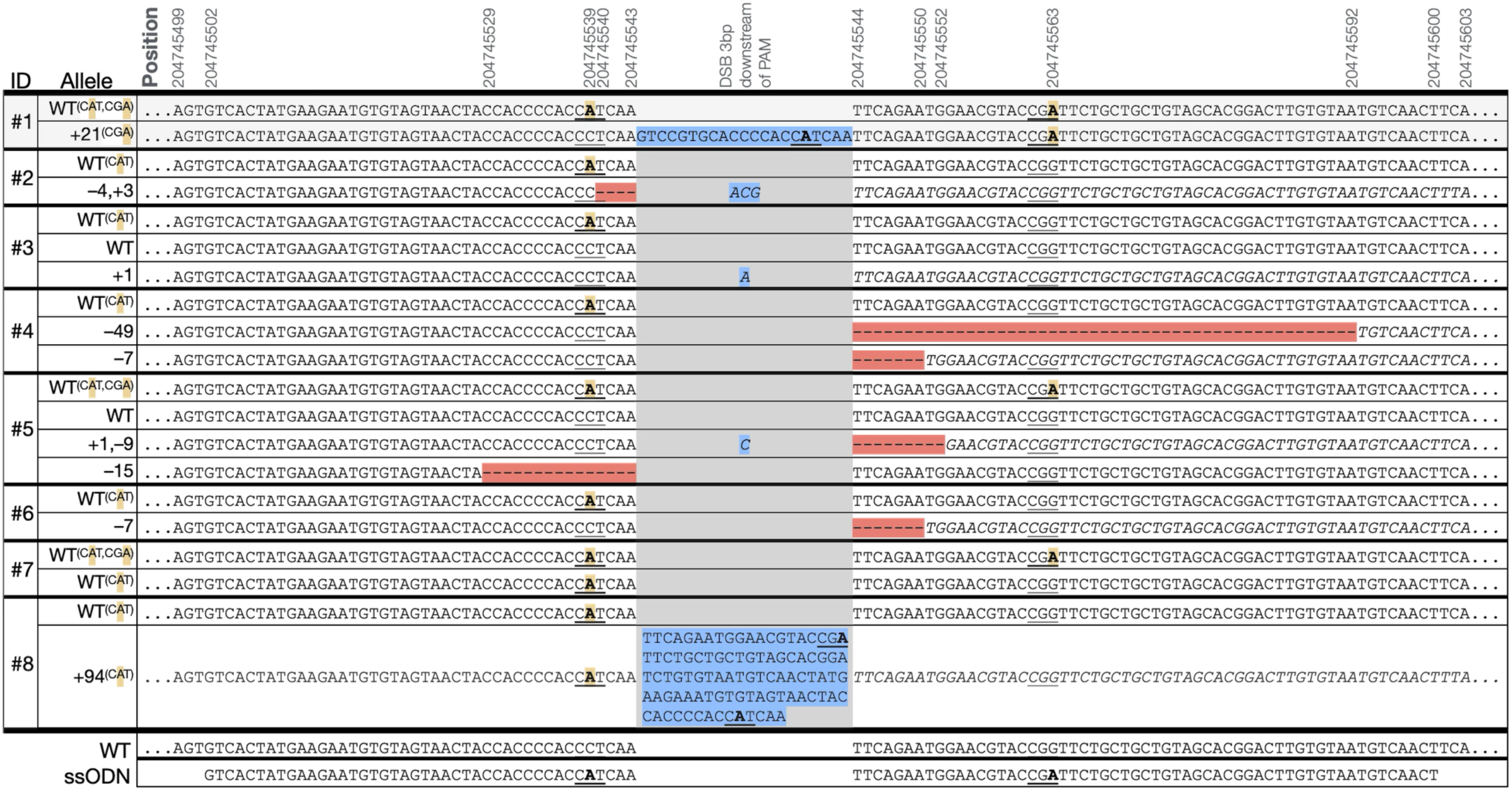
Edited BMPR2 alleles identified by EZ-Amplicon sequencing. Alleles shown were observed in >1% of EZ-Amplicon reads. Positions are on *Ovis aries* Chr2 (NC_056055.1). Red highlighting indicates deleted sequence, blue indicates inserted sequence. protospacer-adjacent motifs (PAMs) are underlined, with disrupting sequences bolded, and highlighted yellow when repaired using the single-stranded oligodeoxyribonucleotides (ssODN) template. The Cas9-induced double stranded break 3 bp downsteam of the PAM is identified in grey, resulting frame-shifted nucleotides are indicated in italics. The additional silent PAM disruption by the ssODN (CGG->CGA) at 204745563 was to disrupt the PAM for sgRNA2; C/T variation at 204745602 is a known SNP (rs404303969).

**Table 1.**
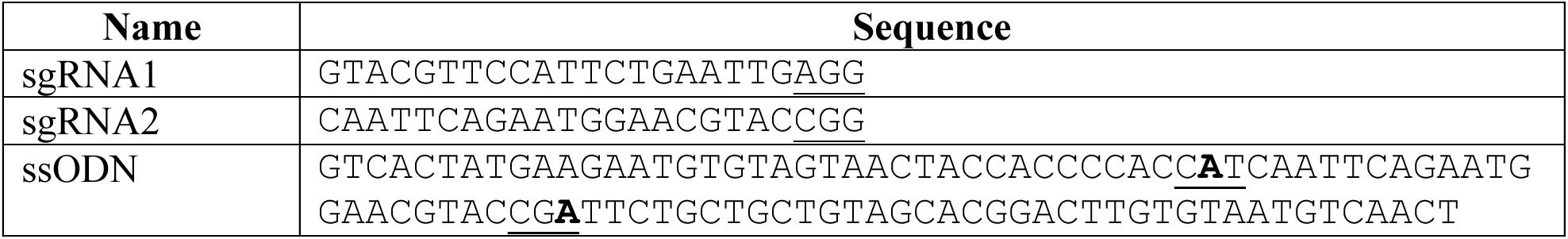
Nucleic acid sequences for blastocyst editing. Intentional mismatches relative to the Oar_v3.1 reference genome are indicated in bold, underlining indicates targeted protospacer-adjacent motifs.

**Table 2.** Genotype characteristics of eight gene-edited lambs. All eight of the gene-edited lambs carried one allele expected to produce a wildtype (WT) protein, edited via homology directed repair using the ssODN template for protospacer-adjacent motif disruption (CCT > CAT). Major allele percentages were determined by tallying the percentage of reads from Illumina EZ-Amplicon sequencing, alleles with sequences represented in less than 1% of total reads were excluded.

| ID | Sex | GT | Status | Sample | Allele 1 | Allele 2 | Allele 3 | Allele 4 |
| --- | --- | --- | --- | --- | --- | --- | --- | --- |
| 1 | F | (+/-) | Perinatal death | Tissue | WT<sup>CAT, CGA</sup> (56%) | +21<sup>CGA</sup> (44%) | N/A | N/A |
| 2 | M | (+/-) | Perinatal death | Tissue | WT<sup>CAT</sup> (49%) | -4,+3 (51%) | N/A | N/A |
| 3 | F | (+/-) | Perinatal death | Tissue | WT<sup>CAT</sup> (35%) | WT (35%) | +1 (30%) | N/A |
| 4 | M | (+/-) | Liveborn | Blood | WT<sup>CAT</sup> (38.5%) | -49 (58%) | -7 (3.5%) | N/A |
|  |  |  |  | Sperm | WT<sup>CAT</sup> (26%) | -49 (58%) | -7 (17%) | N/A |
| 5 | F | (+/-) | Liveborn | Blood | WT<sup>CAT, CGA</sup> (26%) | WT (16%) | +1,-9 (49%) | -15 (8%) |
| 6 | F | (+/-) | Liveborn | Blood | WT<sup>CAT</sup> (52%) | -7 (48%) | N/A | N/A |
| 7 | F | (+/-) | Perinatal death | Tissue | WT<sup>CAT, CGA</sup> (50%) | WT<sup>CAT</sup> (50%) | N/A | N/A |
| 8 | F | (+/-) | Liveborn | Blood | WT<sup>CAT</sup> (78%) | +94<sup>CAT</sup> (22%) | N/A | N/A |

### Expression Analysis

To evaluate the impact of *BMPR2* gene editing on transcript expression, we quantified wildtype *BMPR2* mRNA levels by qPCR in either lung or ear skin from liveborn F0 lambs and normalized expression to the reference gene *ACTB* (Figure 4). *BMPR2* mRNA levels in ear skin were reduced in lamb #4 relative to wildtype (WT) (p=0.003). *BMPR2* mRNA levels in lung tissues were reduced in lamb #5 relative to WT (p=0.006), but the reduction did not reach significance in lamb #6 (p=0.1). The 94 bp insert in exon 3 of *BMPR2* in Lamb #8 reconstituted the *BMPR2* WT sequence such that our qPCR assay for WT *BMPR2* mRNA could not be used for this allele. We therefore extracted RNA from ear skin of lambs #4, #8, and a WT control and did semi-quantitative RT-PCR to visually assess the relative levels of their *BMPR2* transcripts (Figure 5). The WT allele was visible in all three lambs along with the 49 bp deletion allele in lamb #4, which had approximately the same intensity as the lamb #4 WT allele. The 7 bp deletion allele in lamb #4 was indistinguishable from the WT allele. The 94 bp insertion allele in lamb #8 appeared less intense than the lamb #8 WT allele (Figure 5).

**Figure 4.**
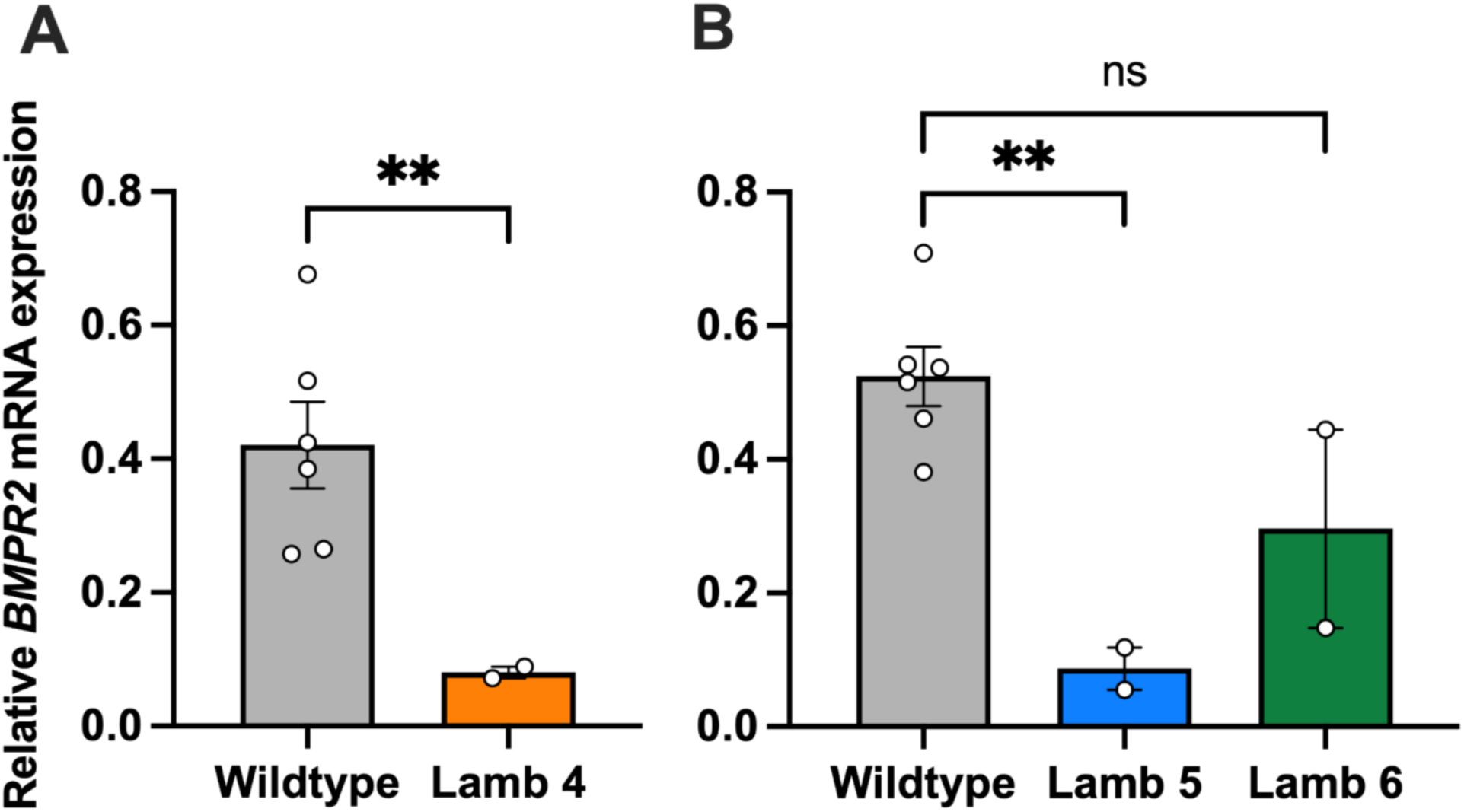
Relative *BMPR2* mRNA expression in wildtype (WT) and *BMPR2*-edited F0 lambs quantified by qPCR. (A) Expression analysis of liveborn lamb #4 ear sample compared with three WT controls, assayed twice. Mean values are shown as bars with individual data points overlaid. (B) Expression analysis of liveborn lamb #5 and #6 lung samples compared with three WT controls assayed twice.

**Figure 5.**
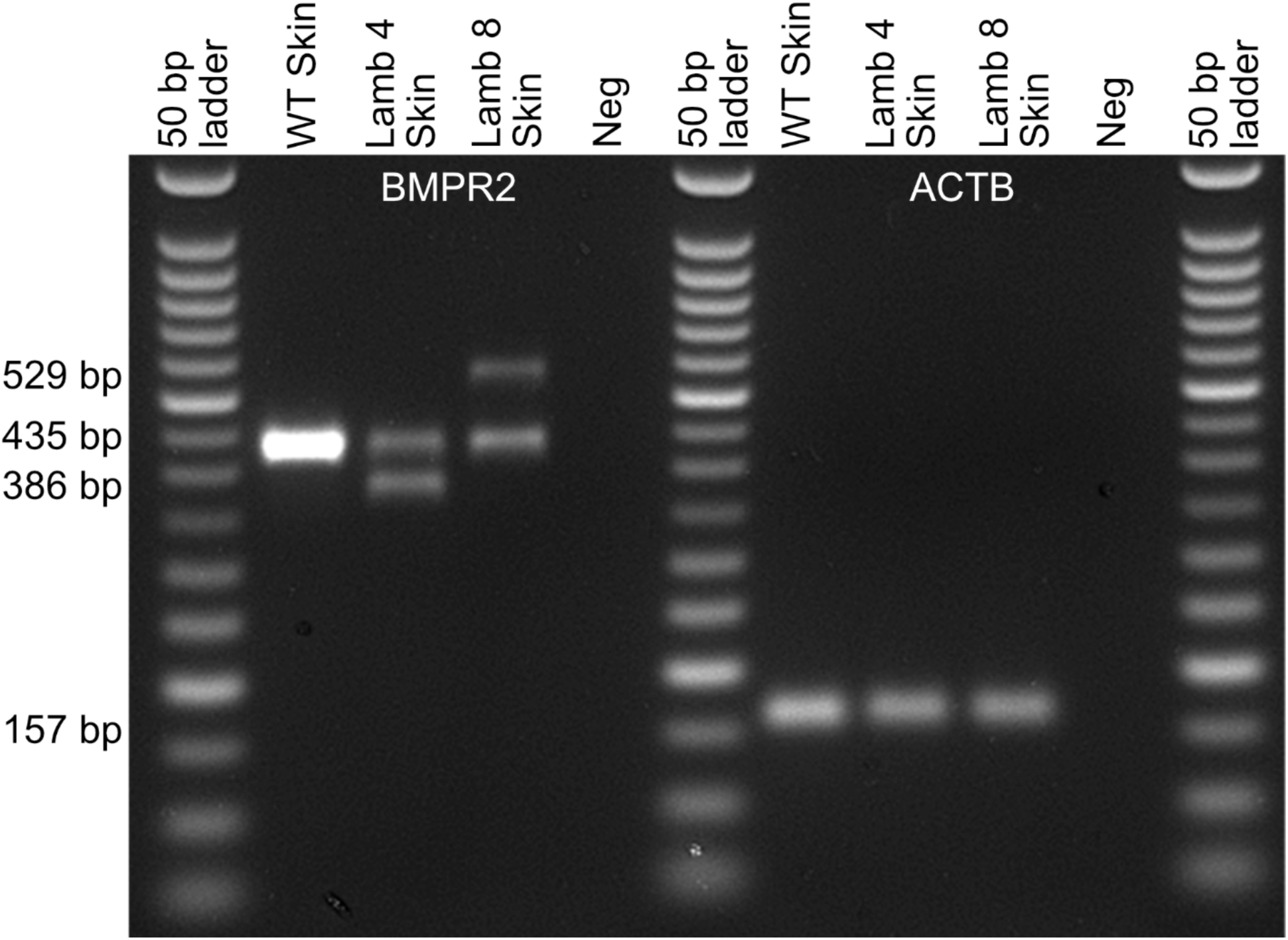
Expression of *BMPR2* transcripts in skin of WT and *BMPR2*-edited F0 lambs #4 and #8. The WT *BMPR2* allele (435 bp), 49 bp deletion allele (386 bp) and 94 bp insertion allele (529 bp) are easily visualized. *ACTB* expression (157 bp) was used as a housekeeping control.

### Cardiac Evaluations

Of the four liveborn gene-edited lambs, three were female. At three weeks of age, echocardiograms were performed on all four *BMPR2* heterozygous lambs. Compared to control (Figure 6A), in two of the three female *BMPR2^(+/−)^*lambs (#6, #8), there was evidence of end systolic septal flattening and right ventricular hypertrophy, with an elevated left ventricular eccentricity index of 1.6 and 1.5, respectively (Table 3). A left ventricular eccentricity index above 1.1 is considered abnormal, with a value above 1.3 highly suggestive of pulmonary hypertension. The echocardiogram and left ventricular eccentricity index of the male (#4) and the third female (#5) *BMPR2^(+/−)^*lambs were normal.

**Figure 6.**
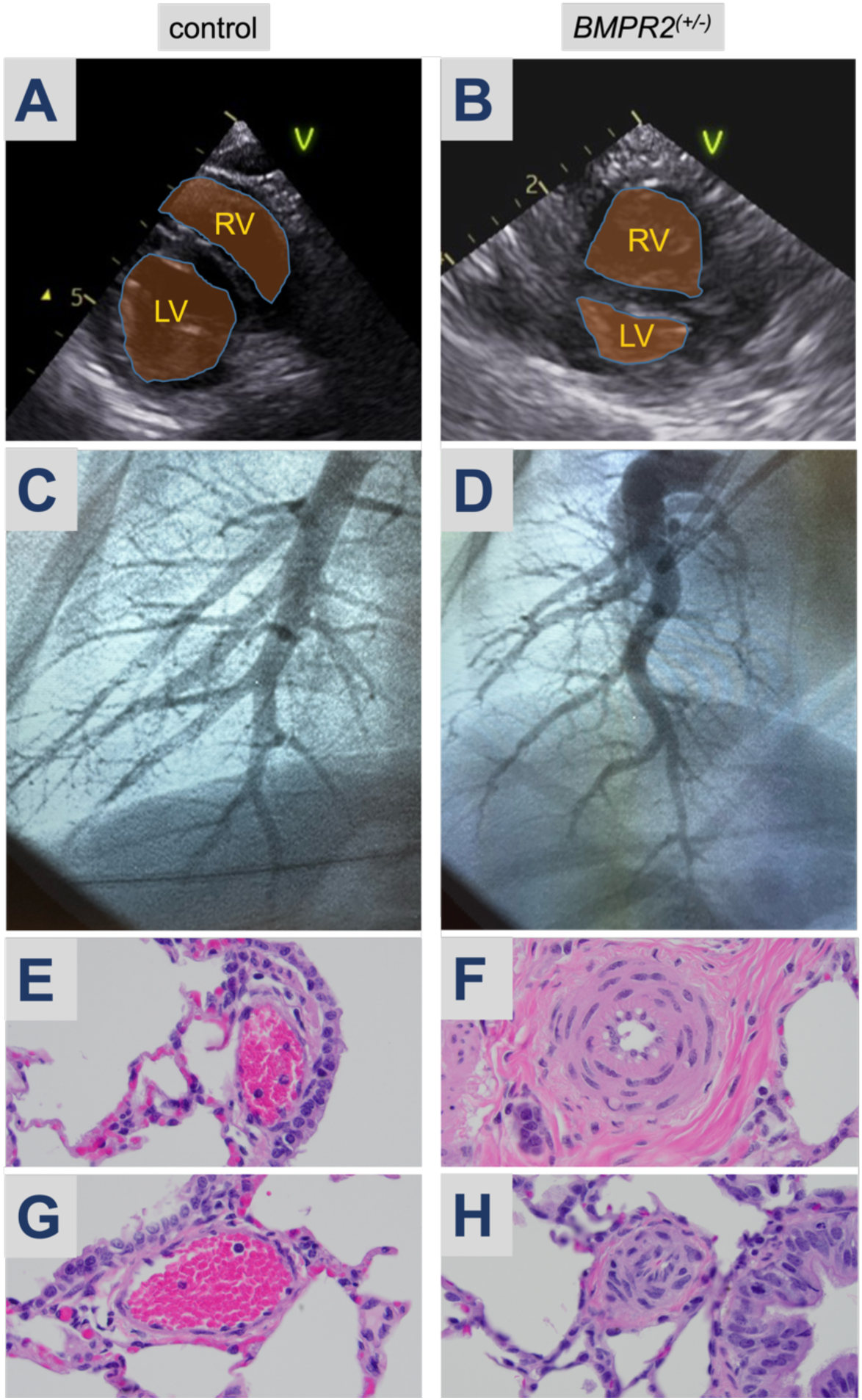
Pulmonary Hypertension Evaluation. Echocardiography at 3-weeks of age, representative images of (A) control versus (B) *BMPR2^(+/−)^* female (#8); RV, right ventricle. LV, left ventricle, V, ventral. (C) Right pulmonary angiography of (C) control and (D) BMPR2*^(+/−)^* lamb (#8). H&E staining at 60x magnification of pulmonary arterioles from *BMPR2^(+/−)^* lambs #5 (F) and #6 (H) associated with medium sized airways. Arterioles from age-matched control lambs (E, G).

**Table 3.**
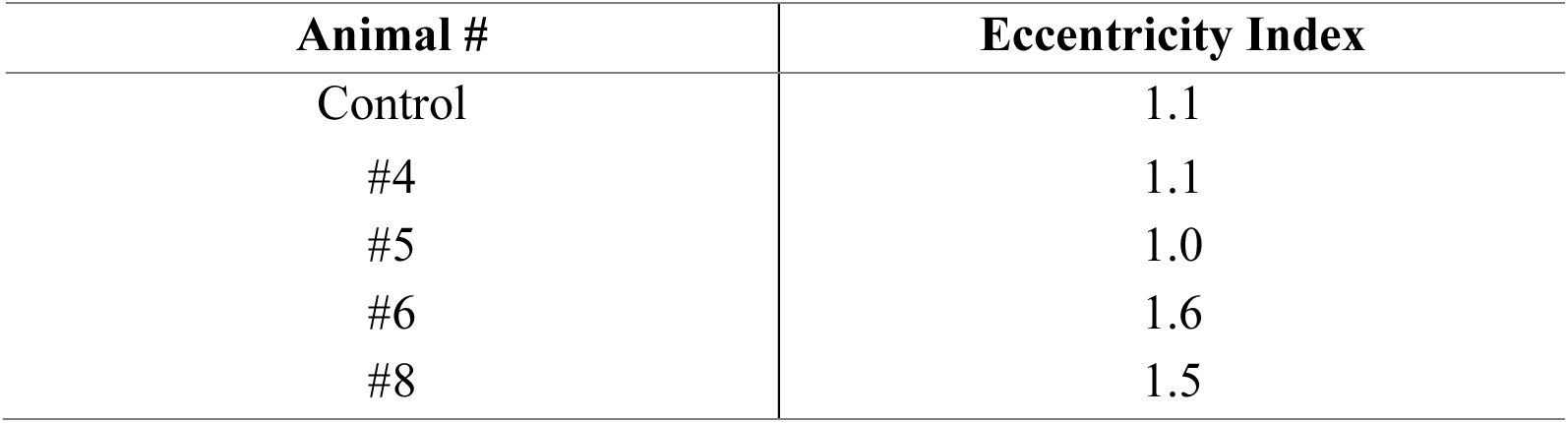
Calculated Eccentricity Index of Left Ventricular Dysfunction in Liveborn *BMPR2^(+/−)^* F0 Lambs versus Control.

| Animal # | Eccentricity Index |
| --- | --- |
| Control | 1.1 |
| #4 | 1.1 |
| #5 | 1.0 |
| #6 | 1.6 |
| #8 | 1.5 |

Each of the three female *BMPR2* heterozygous female lambs were noted to have a persistently patent ductus arteriosus (PDA), at 7 to 8 weeks of age. In contrast, a PDA was not observed in any wildtype lambs by 4-weeks of age, n=6 (by computed tomography angiography at 3 to 4 weeks of age; data not shown).

At 11-weeks of age, we performed a right heart catheterization in *BMPR2^(+/−)^* ewe lamb #8, and an age-matched control (Table 4). In each lamb, there was no PDA. In 21% FiO_2_, the mean pulmonary artery pressure of the *BMPR2^(+/−)^* lamb was 29mmHg, and the calculated resistance (Rp:Rs) was 0.79, both abnormally elevated. In response to acute vasodilator testing (100% O_2_ + 40 ppm inhaled nitric oxide), the *BMPR2^(+/−)^* lamb #8 had a 41% decrease in mean pulmonary artery pressure and a 58% decrease in Rp:Rs. Compared with a normal pulmonary artery morphology in the control lamb (Figure 6D), right pulmonary angiography demonstrated a tortuous and pruned vasculature in the *BMPR2^(+/−)^* lamb that is characteristic of PAH (Figure 6E).

**Table 4.** Hemodynamics from Right Heart Catheterization of a *BMPR2^(+/−)^* F0 Lamb (#8) versus Control. First condition in 21% FiO_2_; second condition in parentheses, in 100% O_2_ + 40ppm inhaled nitric oxide. Qp:Qs, calculated ratio of pulmonary to systemic blood flow; Rp:Rs, calculated ratio of pulmonary to systemic vascular resistance.

| Site | Mean pressure (mmHg) |  |
| --- | --- | --- |
|  | Control<sup>(+/+)</sup> | *BMPR2*<sup>(+/-)</sup> |
| Superior Vena Cava | 5 (4) | 3 (3) |
| Pulmonary Artery | 13 (11) | 29 (17) |
| Pulmonary Artery (wedge) | 6 (7) | 3 (3) |
| Femoral Artery | 65 (60) | 54 (49) |
| Qp:Qs | 1 (1) | 1 (1) |
| Rp:Rs | 0.11 (0.07) | 0.79 (0.33) |

### Lung histopathology

Two of the female F0 *BMPR2^(+/−)^* lambs (#5, #6) were euthanized at 6 and 8 weeks of age, respectively. The first was unable to bear weight on her front leg and thus was unable to stand or ambulate; the second developed sepsis that was non-responsive to treatments. Post-mortem lung samples were processed and compared to an age matched control. In both euthanized *BMPR2^(+/−)^*lambs (Figure 6F, #5; Figure 6H, #6), there was prominent medial hypertrophy with focal occlusive changes of pulmonary arteries associated with medium-sized airways compared with age-matched controls (Figures 6E, 6G). Pentachrome staining highlighted the arterial wall hypertrophy in both female F0 *BMPR2^(+/−)^* lambs, which was more diffuse and severe in #6 than #5 (Figure 7C vs 7B). Smooth muscle actin (SMA) and von Willebrand Factor (vWF) highlighted arterial muscle cell hypertrophy and focal endothelial cell hyperplasia (Figures 7E, 7F). Furthermore, Ki67 staining identified proliferating arterial endothelial (closed arrowheads) and smooth muscle cells (open arrowheads) only in the lung from *BMPR2^(+/−)^* lambs (Figures 7H, 7I).

**Figure 7.**
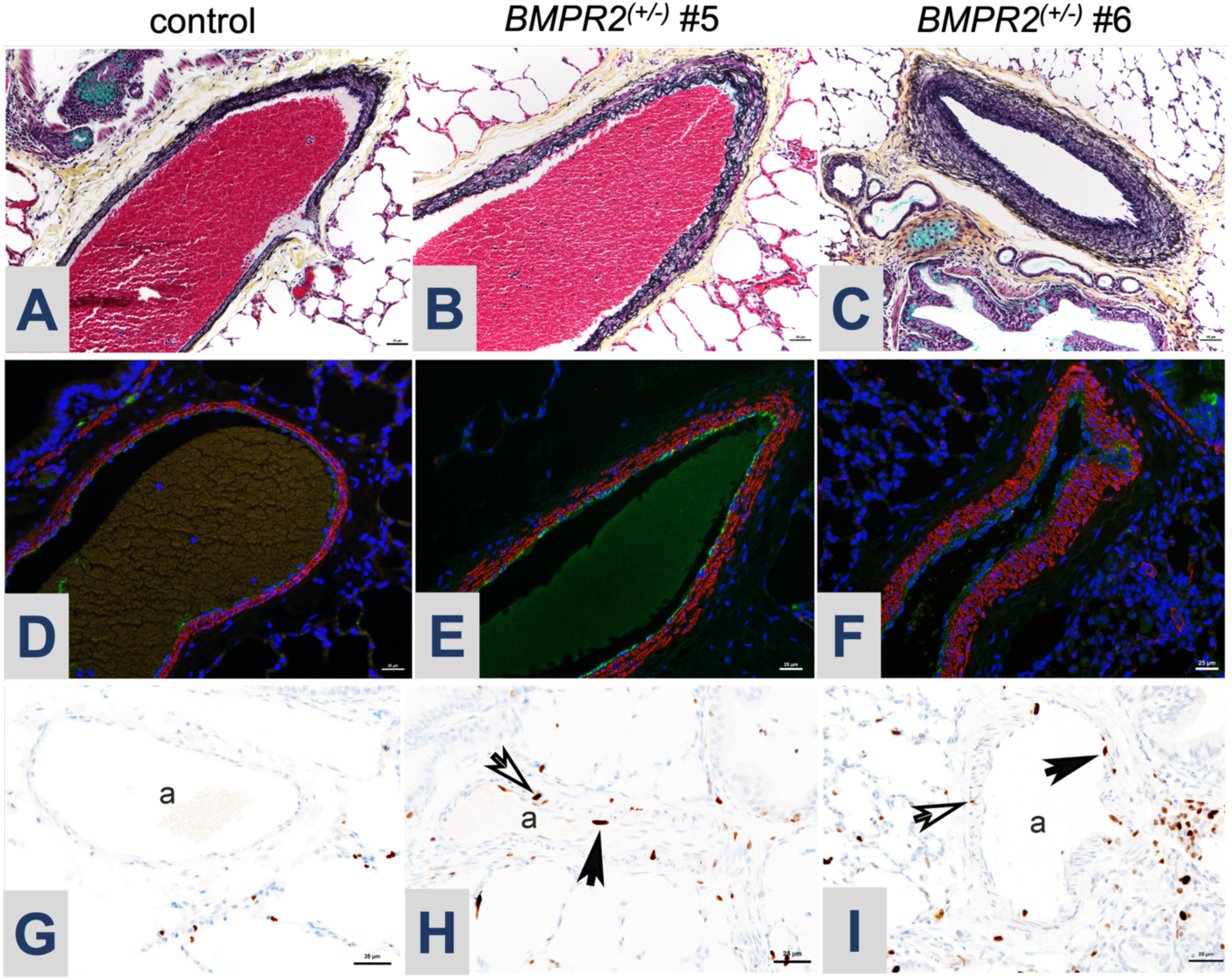
Histopathology of pulmonary arteries from lungs from *BMPR2^(+/−)^* lambs (#5 and #6) compared with control. (A-C) Pentachrome staining identifies arterial hypertrophy in *BMPR2^(+/−)^* lambs; (D-F) Immunofluorescence staining highlights expansion of both endothelial (von Willebrand factor (vWF), green) and smooth muscle (smooth muscle actin (SMA), red) layers. (G-I) Ki67 staining: open arrowheads mark Ki67-positive proliferating cells in the smooth muscle layer and the closed arrowheads mark Ki67-positive proliferating cells in the endothelium; a, pulmonary artery.

### Germline Transmission of Mutant Allele

Illumina EZ Amplicon sequencing of sperm collected from the *BMPR2^(+/–)^* ram lamb #4 revealed a higher relative proportion of a secondary knockout allele in his germline compared to his blood: the 7 bp deletion represented 17% of sperm reads, while the 49 bp deletion was consistent at 58% (Table 2).

Blastocysts created by in vitro fertilization using semen collected from the *BMPR2*^(+/–)^ ram lamb #4 were 47.8% (11/23) WT/-49 bp genotype, 17.4% (4/23) WT/-7 bp genotype, and 34.7% (8/23) were WT/WT^CAT^ with one allele having been generated by HDR repair with the telltale silent “CAT” PAM-disrupting sequence at the sgRNA 1 PAM site (Supplementary Figure 1).

## Discussion

### Heterozygous *BMPR2* Knockout Model Generation

We have successfully developed the first heritable large animal model of pulmonary arterial hypertension. An ssODN was used to produce heterozygous *BMPR2^(+/–)^* knockout sheep by introducing a silent mutation at the PAM site to prevent further editing in one allele while the second allele was disrupted by NHEJ-induced indels. Heterozygous editing was essential to avoid the embryonic lethality associated with complete loss of BMPR2 (12). To our knowledge this is the first documentation of liveborn sheep generated with an intentional CRISPR-Cas9 induced heterozygous edit, resulting in four liveborn gene-edited lambs, each uniquely heterozygous for *BMPR2*.

During CRISPR-Cas9 mediated genome editing, the process of HDR using a ssODN repair template is referred to as single-stranded template repair and results in higher genome editing efficiencies than HDR pathways that use dsDNA repair templates. It has been observed that knock-in rates are higher when the ssODN is complementary to the PAM-containing (non-target) strand (28). This has been linked to the direction of DNA resection after Cas9 cleavage. We found that rates of knock-in were higher with sgRNA1 where the 99 bp ssODN was complementary to the PAM-containing strand than with sgRNA2, which targeted the other strand (Figure 1). In this experiment all eight of the gene-edited lambs showed evidence that at least one allele was repaired using the ssODN template (Table 2). Each contained the synonymous PAM disrupting sequence (CAT) targeted in this experiment (NC_056055.1:g204745539), indicating successful use of the ssODN template. However, the silent substitution on the ssODN (CGA) (NC_056055.1:g204745563), located downstream of the exon 3 cut site, enabled us to determine that the entirety of the 99 bp ssODN was only used in the HDR repair of one wild type allele in three lambs (#1, #5, and #7). It was also notable that in two progeny, #1 and #8, which incorporated 21 bp and 94 bp insertions, respectively, the ssODN template was incorporated into the insertion, as evidenced by the sequence of the PAM-disrupting mutations (Figure 3).

In instances where the double-stranded break was repaired by NHEJ, variations in indel mutations led to the generation of different knockout alleles in most animals. In some instances, multiple alleles were present, suggesting that some editing occurred after the one-cell zygote stage. In the male BMPR2^(+/–)^ lamb (#4), a third allele was present at a higher relative proportion in the germline compared to blood, meaning that he would be expected to produce ∼2/3 *BMPR2^(+/–)^* and 1/3 *BMPR2^(+/+)^* offspring versus the typical 1:1 ratio. This 2 to 1 heterozygous:wildtype *BMPR2* ratio was confirmed by germline testing of his sperm (Supplementary Figure 1).

Each of the eight gene-edited lambs that developed to term had at least one wildtype allele supporting the idea that loss of both *BMPR2* alleles is embryonic lethal. The four lambs that died perinatally were post term and appeared to have died of complications of failed labor associated with large offspring syndrome, a known complication of in vitro fertilization in sheep (29).

### BMPR2 Knockout Model Phenotypic Analyses

By two months of age, each of the three F0 *BMPR2^(+/–)^*female lambs demonstrated evidence of pulmonary vascular disease. Consistent with the variable penetrance and expressivity seen in patients that carry a *BMPR2* mutation, there was a gradation of disease severity. One *BMPR2^(+/−)^*female exhibited a severe PH phenotype by 5 weeks of age, with echocardiographic findings of a globular hypertrophic right ventricle, septal flattening, and an abnormally elevated eccentricity index (Figure 6, Table 3). A subsequent right heart catheterization at 11-weeks of age confirmed significant PAH: she had an elevated mean pulmonary artery pressure greater than 20mmHg (29mmHg in 21% FiO_2_) with a normal pulmonary arterial wedge pressure and an elevated pulmonary vascular resistance (Table 4), as well as a tortuous right pulmonary artery by angiography with evidence of distal arterial pruning.

Each of the two F0 *BMPR2* heterozygous females that were euthanized showed histopathologic evidence of pulmonary arterial involvement, with hyperproliferation and hypertrophy of medium and smaller arterioles in all three vessel layers (Figure 7), and evidence of luminal obliteration (Figure 6F, 6H). Importantly, only one of these lambs (#6) had an abnormal echocardiogram. *BMPR2^(+/−)^* ewe lamb #5 had normal echocardiographic findings and a normal calculated eccentricity index (Table 3), suggesting a disparity between histopathology and echocardiographic evidence of PAH. Interestingly, the male *BMPR2^(+/−)^* lamb (#4) had normal echocardiograms and hemodynamics and was clinically asymptomatic with a histologically normal pulmonary vasculature (data not shown). Critically, two mutant *BMPR2* alleles are represented in his germline (Table 2), and these sperm are functional and can produce viable blastocysts bearing wildtype, −49, or −7 bp alleles (Supplementary Figure 1), which suggests that, if bred to wildtype ewes, this *BMPR2* heterozygous ram should be able to sire *BMPR2^(+/−)^* and wildtype F1 sibling offspring for sustained future studies.

The observation that the three female *BMPR2* heterozygous lambs had a PDA is intriguing. To our knowledge, PDA has not been described as a phenotype in rodent investigations of *Bmpr2* mutations (30–34). Of note, we have never previously observed a PDA in any wildtype lamb that we evaluated. *BMPR2* mutations may be associated with congenital heart defects that include PDA (35), but it is not at all clear that *BMPR2* mutations play a causal role in the development of PDA or other congenital heart disease. Similarly, it is unknown whether delayed closure of the ductus arteriosus is a marker of the severity of pulmonary vascular disease in these female *BMPR2* heterozygous lambs, or might be contributing to their pathophysiology, and warrants further study.

In summary, this new large animal model of heritable PAH offers human-scale assessment of cardiopulmonary physiology, such as the evaluation of right ventricle-pulmonary artery coupling, proximal pulmonary artery mechanics, and pulmonary compliance. In addition, it will allow meaningful investigation into the effects of normal development on disease progression, particularly with respect to the transition to sexual maturity and exposure to clinically relevant ‘second hits’ like pregnancy (36, 37), high altitude (36), and congenital heart disease (38, 39). Lastly, it may allow for preclinical device and drug delivery studies, biomarker discovery, and testing of promising therapeutics, including gene therapies that will lead to improved quality of life for patients affected by PAH.

## Methods

### Sex as a Biological Variable

Our study evaluated both female and male F0 animals, and our findings support what has been observed in humans: variable penetrance of phenotype that is expressed more frequently in females.

### Editing Construct Design

A sgRNA CRISPR/Cas9 approach to target exon 3 of the ovine *BMPR2* gene, in combination with a ssODN, PAM-disrupting, synonymous HDR template was employed to generate heterozygous *BMPR2^(+/–)^* sheep (Figure 1). Two sgRNAs (1 and 2) targeting *BMPR2* exon 3 on opposite strands were designed using *Ovis aries* reference genome Oar_v3.1 (RefSeq GCF_000298735.1) in ChopChop (40), while the 99 bp ssODN was manually designed to silently disrupt the NGG PAM of both guides by inducing a G > T mutation at one of the two Gs in each NGG PAM sequence (Table 1).

### Presumptive Zygote Production

Ovine ovaries were collected fresh from a local abattoir (Superior Farms, CA), stored in 37°C saline and aspirated within 2h of collection. Cumulous Oocyte Complexes (COCs) were collected with a 21 G needle, those with several layers of cumulous cells were matured in batches of 50 for 22-24h in 400 µl of maturation media (BO-IVM, IVF Bioscience, Cornwall, UK) in a humidified incubator at 38.5°C, 5% CO_2_.

Following maturation, COC batches were rinsed 3 times in in vitro fertilization (IVF) media and placed in 400 µl IVF media (BO-IVF, IVF Bioscience). Frozen semen straws, from a Hampshire ram, were thawed in 37°C water for 1 minute prior to washing twice in 1 ml semen preparation media (BO-Semen Prep, IVF Bioscience), centrifuging at 328 *g* for 5 min. Fertilization media was used to adjust sperm concentration to 2 × 10^6^ sperm/ml, 50 µl of which was used to fertilize each batch for 6 h in a humidified incubator at 38.5°C, 5% CO_2_.

### Electroporation and Embryo Culture

Presumptive zygotes 6 h post-fertilization were denuded by 3 minutes of vortexing in SOF-HEPES, then washed 3x in SOF-HEPES and 3x in Opti-MEM (Thermo Fisher Scientific, Waltham, MA). Zygotes were moved into a fresh drop of Opti-MEM and mixed with the ssODN alongside ribonucleoprotein complexes comprised of 100 ng/µl sgRNA 1 or 2 and 200 ng/µl Cas9 and 600 ng/ul ssODN in a 20 µl final volume. The mixture was loaded into a 1 mm Electroporation Cuvette (Bulldog-Bio, Portsmouth, NH) and electroporated via NEPA21 Super Electroporator using poring conditions of 2 bipolar pulses of 30 volts for 3.5 msec and transfer conditions of 50 msec each. Recovered presumptive zygotes were washed 3x in SOF-HEPES and groups of 50 were placed in 500 µl of culture media (BO-IVC, IVF Bioscience) covered with 400 µl mineral oil. Embryos were cultured for 7 days in a humidified incubator at 38.5°C, 5% CO_2_, 5% O_2_, and 90% N_2_.

### PCR Analyses

For brevity, conditions (primer sequences, reagent composition, thermocycling conditions) for all PCR assays in subsequent sections are presented in Supplementary Table 1.

### Genomic Analyses of Blastocysts

At seven days, zygotes that had advanced to the blastocyst stage of development were collected in 10 µl QuickExtract^TM^ DNA Extraction Solution (Biosearch Technologies), and incubated at 65°C for 6 mins, 98°C for 2 mins, then cooled to 4°C to collect DNA. PCR products were gel-purified prior to submission for Sanger sequencing (Genewiz, South San Francisco, CA). Sequences were analyzed using TIDER (41), a computer model that quantifies the frequency of targeted small nucleotide changes introduced by CRISPR in combination with HDR using a donor template.

### Embryo Transfer

Eazi Breed™ CIDRs® Progesterone inserts (Zoetis, Parsippany, NJ) were placed for 6 days to introduce estrous synchronize multiparous surrogate ewes at insert removal intramuscular injection of prostaglandin F2-alpha analog (10 mg dinoprost thrometamine; Zoetis) and PG600 (400 IU PMSG, 200 IU hCG; Intervet, Madison, NJ) was given. Ewes were food-restricted for 24h and water-restricted 12h before undergoing a minimally invasive laparoscopic procedure, as previously described (42). Briefly, ewes were sedated by intravenous injection of 1.1-2.2 mg/kg of ketamine and 0.2-0.3 mg/kg of Midazolam prior to surgery. 2% Lidocaine was used as a local anesthetic at the incision site. Ewes were placed in the Trendelenburg position and had their abdomen insufflated with CO_2_. The ovaries and uterus were visualized via 0-degree laparoscope in the caudal abdomen, and 5-6 blastocysts were transferred into the uterine horn ipsilateral to the corpus luteum.

### Genotype Analysis of F0 Lambs

Genomic DNA was extracted from either tissue from stillborn lambs or blood from liveborn lambs using the DNeasy Blood and Tissue Kit (Qiagen, Redwood City, CA), PCR products were gel purified (Figure 2), including two bands extracted independently for both lambs #4 and #8, and subjected to Sanger sequencing (Genewiz). For quantification of alleles and identification of rare alleles in mosaic lambs DNA was PCR amplified, all products 300-500 bp were gel purified as a single sample per lamb and subjected to Amplicon-EZ Next Generation Sequencing (Genewiz).

### RNA Extraction and RT-PCR

For wildtype (WT) controls, ear, lung and heart tissues were collected under hygienic conditions from sheep immediately following slaughter at an abattoir, snap frozen in liquid nitrogen, and stored at –80°C until processing. Ear tissues from Lambs 4 and 8 were collected by punching the ear of the living animal while lung tissues from Lambs 5 and 6 were collected immediately following euthanasia, snap frozen with liquid nitrogen and stored at –80°C. Total RNA was extracted by homogenizing in Qiazol reagent (Qiagen) following the manufacturer’s instructions. RNA concentration and purity were determined using a NanoDrop1000^TM^ (Thermo Fisher Scientific, USA). RNA integrity was verified by confirming presence of non-degraded 28S and 18S bands on a MOPS-formaldehyde 1.2% agarose gel electrophoresis. Total RNA was treated with TURBO DNase^TM^ (Invitrogen, Carlsbad, CA) according to the manufacturer’s instructions. For cDNA synthesis, 1 µg of total RNA was reverse transcribed using GoScript^TM^ Reverse Transcriptase in 20 µL reactions according to manufacturer’s instructions with included Oligo(dT) (Promega), supplemented with 50 ng of random hexamers. Each RT reaction was accompanied by no RT control in which the reverse transcriptase was replaced by RNase free water, was validated by PCR. cDNA was diluted with an equal volume of RNase free water and stored at −80°C.

Expression of BMPR2 alleles in ear skin tissue from a WT sheep, and from Lambs 4 and 8 was examined by semi-quantitative PCR on cDNA, resulting PCR products were run on a 2% agarose gel where the size and intensity of the bands were estimated.

### Quantitative PCR (qPCR)

Beta actin (*ACTB*) was used as housekeeping control and primers for *BMPR2* and *ACTB* Taqman quantitative PCR (qPCR) reactions were designed to span exon–exon junctions, when possible, to avoid genomic DNA amplification (Supplementary Table 1). Locking Nucleic Acid Probes for WT *BMPR2* that included either the WT CCT sequence, or the sgRNA 1 PAM-disrupting “CAT” sequence encoded in the ssODN (Figure 1) were designed using OligoArchitectTM Online (Sigma-Aldrich, St. Louis, MO). Probes were labeled 3’ with black hole quencher-1 (BHQ-1) and 5’ with either 6-carboxyfluorescein (6-FAM) or SUN (Integrated DNA Technologies, Coralville, Iowa). Each PCR run included a no-template control containing all reagents except cDNA. All samples, standards and controls were assayed in duplicate.

The level of BMPR2 and ACTB mRNA expression were determined using a relative standard curve prepared from six four-fold serial dilutions of a lung RNA sample known to have high levels of BMPR2 gene expression. A standard curve was used on every qPCR plate. All standard curves had a linear regression coefficient of determination of at least 97 and ACTB’s efficiency was 86% while BMPR2’s was 77%. Standard curves were generated by linear regression using Ct versus log (dilution factor). The BMPR2 and ACTB levels in each sample were calculated from Ct values using the standard curve. Data were expressed as the ratio between BMPR2 and ACTB expression levels, yielding a normalized relative expression level of BMPR2 mRNA.

### Germline Transmission by *BMPR2^(+/–)^* F0 Ram

Semen was collected from the gene-edited ram (#4) following sexual maturity. DNA was extracted from the semen, BMPR2 was PCR amplified and submitted for Amplicon-EZ Next Generation Sequencing (Genewiz). The semen was also used to fertilize sheep oocytes *in vitro* as described previously. The resulting blastocysts were analyzed as described (Genomic Analyses of Blastocysts) and PCR products were sequenced using Sanger sequencing to identify which allele was transmitted.

### Echocardiograms of *BMPR2^(+/–)^* F0 Lambs

Transthoracic echocardiograms were performed on non-sedated 3-week-old lambs by a single echocardiographer. The coat was clipped bilaterally and prepared with alcohol and ultrasound gel. Cine-loops were obtained with an ultrasound machine (Vivid IQ Premium V204; GE Healthcare, Chicago, IL) equipped with a 6S-RS Sector Transducer and simultaneous ECG recording. Standard 2-dimensional views obtained from the right chest wall included 4-chamber apical view, long axis views of the right and left ventricular outflow tracts, and short axis views at the aortic, mitral, and chordal level. M-mode images of the aortic valve, mitral valve, and left ventricle at the chordal level were also obtained. From the left chest wall, long axis 2-dimensional images of the left atrium, aortic valve, and pulmonic valve were obtained. A second echocardiographer, who was blinded to the animal identification, evaluated the images in Syngo (Siemens Healthineers, Version VA40D), and using the parasternal short axis views in systole, calculated the eccentricity index as previously described (43).

### Right heart catheterization of *BMPR2^(+/–)^* F0 Lamb

At 11-weeks of age, one female gene-edited lamb (#8) and an age-matched control underwent hemodynamic right heart catheterization via percutaneous access of the right internal jugular vein, using standard evaluation techniques for pediatric pulmonary hypertension (44). Lambs were maintained in a supine position under inhaled anesthesia with 1-3% Isoflurane and were mechanically ventilated using a GE Datex Ohmeda Aestiva5 ventilator. ETCO2 was maintained between 35-45mmHg, with confirmation as needed by arterial blood gases (ABL 90 Flex; Radiometer America, Brea, CA) from a peripheral arterial line. Maintenance fluids (Lactated Ringers) were delivered intravenously during the procedure. Percutaneous internal jugular vein 5 Fr or 6 Fr introducer sheaths were placed, and lambs were heparinized with 150-300 IU/kg Heparin sodium. Activated clotting times (ACTs) were monitored to maintain adequate anti-coagulation (ACT monitor; Actalyke Mini-II; Helena Laboratories, Beaumont, TX) during the study.

Baseline hemodynamics, pulmonary to systemic blood flow ratio (Qp:Qs), and Fick-derived cardiac index were assessed using a 5 Fr or 6 Fr balloon wedge end-hole catheter under fluoroscopic guidance. The catheterization was performed in multiple conditions: in 21% FiO_2_, with 100% FiO_2_, and with 100% FiO_2_ + 40ppm inhaled nitric oxide (INOmax; Mallinckrodt, St. Louis, MO). VO_2_ and other assumptions by weight and age for ovine models were based on previously published studies of cardiac catheterization in lambs (45–48). Right pulmonary artery angiography was performed with hand injection of radiopaque contrast (Omnipaque 350; GE Healthcare) under fluoroscopy (2005 Powermobil; Siemens, Washington, DC), using techniques as described for clinical practice (49, 50). At the conclusion of the cardiac catheterization, the catheters and percutaneous sheath were removed, manual pressure was held until hemostasis, and the skin was closed with tissue adhesive. The lambs were recovered and monitored until back to baseline activity.

### Histology of *BMPR2^(+/–)^* F0 Lambs

During terminal study, when anesthetized and supported by mechanical ventilation, baseline lung samples were taken from the right middle lobe and immediately flash frozen in liquid nitrogen, placed in phosphate buffered saline with 4% paraformaldehyde, or formalin-fixed and paraffin embedded. Special staining and immunostaining were performed on 5-µm sections of representative lung samples from 1 control and 2 *BMPR2^(+/−)^* lambs. Movat pentachrome stain was performed according to established protocols (51). Immunostaining for Ki67 (RRID:AB_2631211, 1:100; Agilent, Santa Clara, CA) was performed on the Ventana BenchMark Ultra after CC1 antigen retrieval. Dual immunofluorescence for Von Willebrand Factor (VWF; RRID:AB_2315602, 1:100; Agilent) and smooth muscle actin (SMA; RRID:AB_2223500, 1:400; Agilent) was carried out following citrate pH 6.0 antigen retrieval and serum block; antibodies were incubated overnight at room temperature. VWF was detected with donkey anti-rabbit Alexa Fluor 488 and SMA with donkey anti-mouse cyanine (both 1:1000; Jackson ImmunoResearch, West Grove, PA). Coverslips were mounted using Vectashield fluorescent mounting medium with DAPI (Vector Laboratories, Newark, CA). Images were visualized and captured with a digital camera mounted on a Nikon Eclipse 80i microscope using NIS-Elements Advanced Research Software v4.13 (Nikon Instruments Inc., Melville, NY).

### Statistics

A Mann-Whitney test (Prism 10; GraphPad Software, Boston, MA) was used to compare the efficiency of sgRNA 1 versus sgRNA2. Statistical analyses of the qPCR expression data were performed in R (version 2024.09.0 Build 375; 52). For comparisons between two groups (e.g., Lamb #4 vs. wildtype) an unpaired two-tailed Welch’s t test was used. For comparisons among three groups (wildtype, Lamb #5, and Lamb #6), one-way ANOVA was performed followed by a Dunnett’s multiple comparisons test to identify differences to the wildtype. Normality and homogeneity of variance were assessed prior to testing. Results are reported as mean ± SEM, and significance was set at p < 0.05.

### Study Approval

All protocols and procedures related to the care and evaluation of the animals in this study were approved by the Institutional Animal Care and Use Committees (IACUC) of the University of California, Davis and the University of California, San Francisco.

## AUTHOR CONTRIBUTIONS

SAD and ALV designed and coordinated the studies and wrote and edited the manuscript; ARB, DSF, and TFB performed experimental work to optimize editing; BM and TU performed the laproscopic procedures for embryo transfer; OF performed expression analysis; NAW and JFT performed molecular analysis, and wrote and edited the manuscript; JMM performed the echocardiography; HN analyzed the echocardiograms; EKA performed the right heart catheterizations; EGJ performed computed tomography, GHD and OAGV performed lung histology; JRF and EDA designed the studies and edited the manuscript; RH managed all aspects of animal care and their studies, including delivery of the F0 lambs.

## ACKNOWLEDGEMENTS

We are grateful to Christian Vento, Hadiya Manzoor, and Amy Lesneski and her team for their careful animal husbandry and technical assistance. We wish to thank Superior Farms for their donation of sheep ovaries.

