## Supplemental Figure and Tables for "A Large Animal Model of Heritable Pulmonary Arterial Hypertension Using Gene-edited *BMPR2* Sheep"

### Supplementary Figures and Tables

**Supplementary Figure 1.** Analysis of 23 sheep blastocysts produced through in vitro production with Lamb #4 sperm and oocytes derived from slaughterhouse-collected wild type (WT) ovaries. (A) Nested PCR targeting *BMPR2* was performed to identify 11/23 (47.8%) embryos with the -49 bp deletion. (B) Embryos with only the WT-sized band were Sanger sequenced to identify the 4/23 (17.4%) embryos (underlined number) with both WT and the 7 bp deletion alleles. Vertical dashed line indicates canonical cut site of Cas9 with single guide RNA 1.

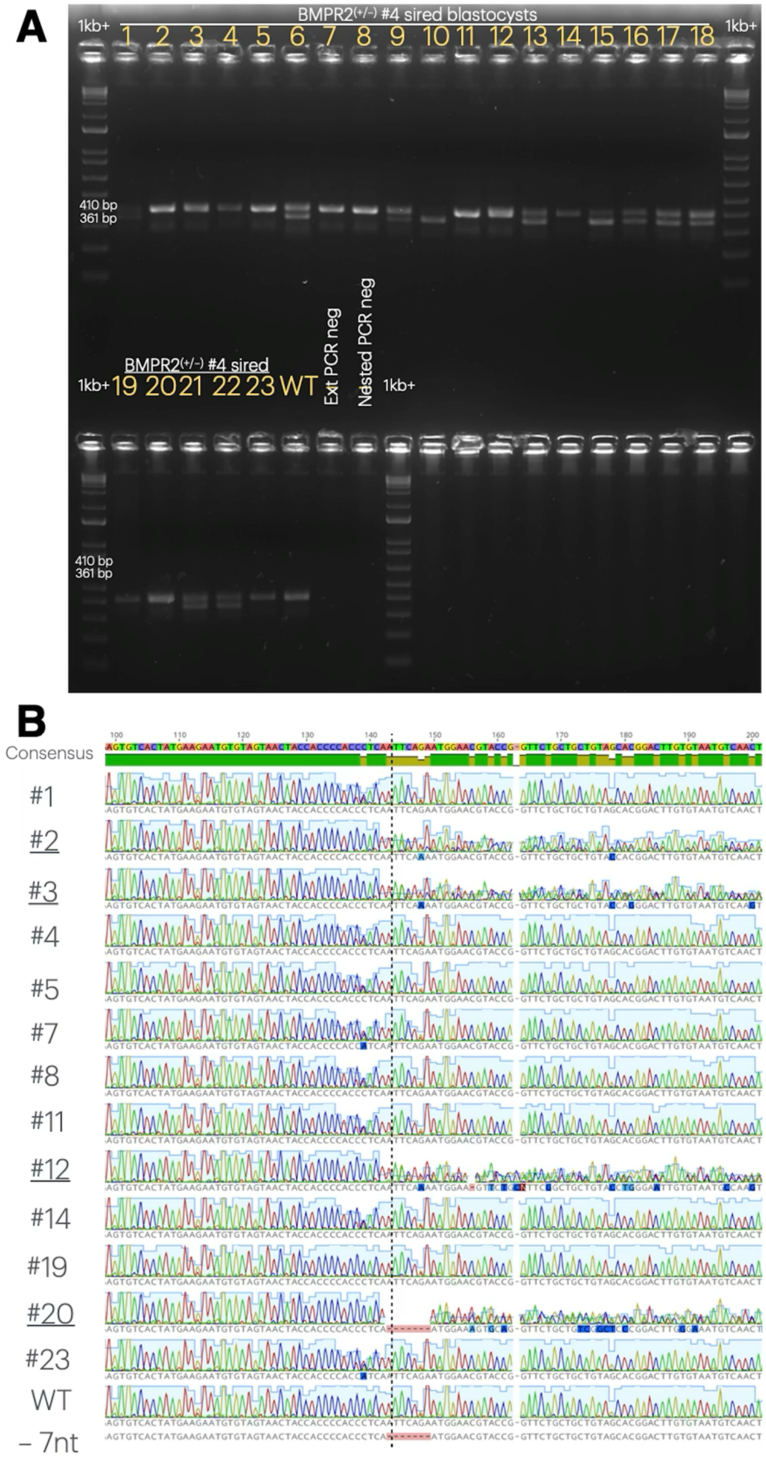

**Supplementary Table 1.** PCR primers, reaction conditions, cycling conditions, expected wild type (WT) product size and quantitative PCR amplification efficiency (E). Primers were 10 µM except where indicated. GoTaq Green, Promega. KAPAG2, Roche. Q5, NEB. Apex Taq, Genesee Scientific.

| Purpose | Primer | Reagents/rxn | Rxn conditions | WT Amplicon Size (bp) | E (%) |
|---|---|---|---|---|---|
| Nested_1 for Blastocysts Sanger Sequencing/ TIDER | Intron2-F: ACAGCAGAAGGACTTAGCCA<br>Intron3-R: TCTTGGAAATCCTGGTGTACCTT | 1 µL Primer Mix<br>10 µL KAPAG2 Robust ReadyMix<br>4 µL Water<br>5 µL Lysis product | 95ºC 5 mins<br>[95ºC 15 s<br>58ºC 30 s<br>72ºC 1 min] x35<br>72ºC 5 mins | 838 | N/A |
| Nested_2 for Blastocysts Sanger Sequencing/ TIDER | Intron2-F2: TTGTTTGCAATTTGTTTGTTGTTTTCC<br>Intron3-R2: TCATAATGCTGGAACTCTCTCTCTT | 1 µL Primer Mix<br>10 µL KAPAG2 Robust ReadyMix<br>9 µL Water<br>5 µL Nested_1 Product | 95ºC 3 mins<br>[95ºC 15 s<br>60ºC 30 s<br>72ºC 1 min] x35<br>72ºC 5 mins | 376 | N/A |
| Genotype Tissues/ Blood | Intron2-F2: TTGTTTGCAATTTGTTTGTTGTTTTCC<br>Intron3-R: TCATAATGCTGGAACTCTCTCTCTT | 1 µL F Primer<br>1 µL R Primer<br>12.5 µL GoTaq Green<br>10.5 µL Water<br>1 µL DNA | 95ºC 5 mins<br>[95ºC 15 s<br>60ºC 30 s<br>72ºC 45 s] x35<br>72ºC 5 mins | 376 | N/A |
| Rare/mosaic allele identification (Lambs 1-7) | Intron2-F2: TTGTTTGCAATTTGTTTGTTGTTTTCC<br>Intron3-R: TCTTGGAAATCCTGGTGTACCTT | 500 nM ea. primer<br>Q5 High-Fidelity Master Mix | 98ºC 30 s<br>[98ºC 10 s<br>60ºC 30 s<br>72ºC 30 s] x35<br>72ºC 5 mins | 431 | N/A |
| Rare/mosaic allele identification (Lamb 8) | Exon3-F: GGTGATCCCCAGGAGTGTC<br>7DelIntron3_R: TCTTTGGAGAAAGGAATTTTCAGGA | 500 nM ea. primer<br>Q5 High-Fidelity Master Mix | 98ºC 2 mins<br>[98ºC 10 s<br>60ºC 30 s<br>72ºC 25 s] x40<br>72ºC 10 mins | 201 | N/A |
| Semi-Quantitiative RT-PCR of BMPR2 (Lambs 4, 8) | cDNAEx2_F: CGTCTCTGTGCATTTAAAGATCCA<br>cDNAEx4_R: TGTCAGCATCCTATATCCAAAGCA | 400 nM ea primer<br>Apex Taq RED | 95ºC 2 mins<br>[95ºC 15 s<br>60ºC 30 s<br>72ºC 45 s] x37<br>72ºC 5 mins | 435 | N/A |
| Semi-Quantitiative RT-PCR of ACTB (Lambs 4, 8) | ACTB_F3: GACACCGCAACCAGTTCG<br>ACTB_R3: CCCACCATCACGCCCTG | 400 nM ea primer<br>Apex Taq RED | 95ºC 2 mins<br>[95ºC 15 s<br>60ºC 30 s<br>72ºC 30 s] x40<br>72ºC 5 mins | 157 | N/A |
| Quantitative PCR | ACTB_qPCR_F: TCCTGGGCATGGAATCCTG<br>ACTB qPCR R: GGGGCGCGATGATCTTGAT<br>ACTB Ex5Probe: 6-FAM/CGGGACCAC/ZEN/CATGTACCCTGGCATCG/IABkFQ<br>BMPR2cDNA_F4: TCCCCAGGAGTGTCACTATGAA<br>BMPR2qPCR R3-2: ATGATTATTGTCTCATCTCGGTTAAAT<br>BMPR2_Ex3_WTC_Probe: SUN/CACC+CTC+AAT+TCA+GAATGG/IABkFQ<br>BMPR2_Ex3_WTA_LNA_Probe: SUN/CACCAT+CAA+TTC+AGA+ATGG/IABkFQ | 5 µL TaqMan Fast Advanced (ABI)<br>900 nM ea. primer<br>150 nM ea. BMPR2 probe<br>250 nM ACTB probe<br>1 µL cDNA<br>Water to 10 µL final vol. | 50ºC 2 mins<br>95ºC 20 s<br>[95ºC 5 s<br>60ºC 20 s] x40 | ACTB: 201<br>BMPR2: 189 | ACTB: 104<br>BMPR2: 97 |
| Nested_1 for Germline Transmission | Intron2-F: ACAGCAGAAGGACTTAGCCA<br>Intron3-R: TCTTGGAAATCCTGGTGTACCTT | 0.4 µL F Primer<br>0.4 µL R Primer<br>10 µL GoTaq Green<br>4.2 µL Water<br>5 µL Lysis product | 95ºC 5 mins<br>[95ºC 15 s<br>55ºC 30 s<br>72ºC 90 s] x40<br>72ºC 5 mins | | |
| Nested_2 for Germline Transmission | oBPMR2-Ex3-NestR2_F: TGTACTCCATCACATTGTTTGCA<br>oBPMR2-Ex3-NestR2_R: GCCATTAGCTAGATGTACTCTCA | 0.4 µL F Primer<br>0.4 µL R Primer<br>10 µL GoTaq Green<br>7.2 µL Water<br>2 µLNested_1 product | 95ºC 2 mins<br>[95ºC 15 s<br>62ºC 30 s<br>72ºC 1 min] x35<br>72ºC 5 mins | 410 | N/A |

**Supplementary Table 2.** Predicted nuclei acid sequences of edited and wild type (WT) BMPR2 alleles. Snapgene software (www.snapgene.com) predicted the amino acid sequences of BMPR2 edits described in Table 2 and Figure 3. *indicates stop codon, in-frame insertions are bolded, frame-shifted amino acids are underlined, in-frame deletion locations indicated by /.

| | |
|---|---|
| **WT and WT (CAT)** | MTSSPRRPRRVPSLLWTVLLVSAAAASQNQERLCAFKDPYQQDLGIGESRISHENGTILCSKGSTC<br>YGLWEKSKGDINLVKQGCWSHIGDPQECHYEECVVTTTPPSIQNGTYRFCCCSTDLCNVNFTENFP<br>PPDTTPLSPPHSFNRDETIIIALASVSVLAVLIVALCFGYRMLTGDRKQGLHSMNMMEAAASEPSL<br>DLDNLKLLELIGRGRYGAVYKGSLDERPVAVKVFSXANRQNFINEKNIYRVPLMEHDNIARFIVGD<br>ERVTADGRMEYLLVMEYYPNGSLCKYLSLHTSDWVSSCRLAHSVTRGLAYLHTELPRGDHYKPAIS<br>HRDLNSRNVLVKNDGTCVISDFGLSMKLTGNXLVRPGEEDNAAISEVGTIRYMAPEVLEGAVNLRD<br>CESALKQVDMYALGLIYWEIFMRCTDLFPGXSVPEYQMAFQTEVGNHPTFEDMQVLVSREKQRPKF<br>PEAWKENSLAVRSLKETIEDCWDXDAEARLTAQCAEERMAELMMIWERNKSVSPTVNPMSTAMQNE<br>RNLSHNRRVPKIGPYPDYSSSSYIEDSIHHTDSIVKNISSEHSMSSTPLTIGEKNRNSINYERQQA<br>QARIPSPETSVTSLSTNTTTTNTTGLTPSTGMTTISEVPYPDETSLHATNVSQPVGPTPVCLQLTE<br>EDLETNKLDPKEVDKNLKESSDENLMEHSLKQFSGPDPLSSTSSSLPYPLIKLAVEVTGQQDFTQA<br>ANGQACLIPDVPPTQIYPLPKQQNLPKRPTSLPLNTKNSTKEPRLKFGSKHKSNLKQVETGVAKMN<br>TINAAEPHIVTVTMNGVAGRNHSVNSHTATTQYANGVVPSGQTANTVAHRAQEMLQNQFIGEDTRL<br>NINSSPDEHEPLLRREQQAGHDEGVLDRLVDRRERPLEGGRTNSNNNNSNPCSEQEVPTQGVPSTV<br>ADPGPSKPRRAQRPNSLDLSATNVLDGSSLQLGDSTQDGKSGSGEKIKKRVKTPYSLKRWRPSTWV<br>ISTEPLDCEVNNNGKDRAVHSKSSTTVYLADGGTATTMVSKDIGMNCL* |
| **#1 Allele 2 (+21)** | MTSSPRRPRRVPSLLWTVLLVSAAAASQNQERLCAFKDPYQQDLGIGESRISHENGTILCSKGSTC<br>YGLWEKSKGDINLVKQGCWSHIGDPQECHYEECVVTTTPPS**SPCTPPS**IQNGTYRFCCCSTDLCNV<br>NFTENFPPPDTTPLSPPHSFNRDETIIIALASVSVLAVLIVALCFGYRMLTGDRKQGLHSMNMMEA<br>AASEPSLDLDNLKLLELIGRGRYGAVYKGSLDERPVAVKVFSXANRQNFINEKNIYRVPLMEHDNI<br>ARFIVGDERVTADGRMEYLLVMEYYPNGSLCKYLSLHTSDWVSSCRLAHSVTRGLAYLHTELPRGD<br>HYKPAISHRDLNSRNVLVKNDGTCVISDFGLSMKLTGNXLVRPGEEDNAAISEVGTIRYMAPEVLE<br>GAVNLRDCESALKQVDMYALGLIYWEIFMRCTDLFPGXSVPEYQMAFQTEVGNHPTFEDMQVLVSR<br>EKQRPKFPEAWKENSLAVRSLKETIEDCWDXDAEARLTAQCAEERMAELMMIWERNKSVSPTVNPM<br>STAMQNERNLSHNRRVPKIGPYPDYSSSSYIEDSIHHTDSIVKNISSEHSMSSTPLTIGEKNRNSI<br>NYERQQAQARIPSPETSVTSLSTNTTTTNTTGLTPSTGMTTISEVPYPDETSLHATNVSQPVGPTP<br>VCLQLTEEDLETNKLDPKEVDKNLKESSDENLMEHSLKQFSGPDPLSSTSSSLPYPLIKLAVEVTG<br>QQDFTQAANGQACLIPDVPPTQIYPLPKQQNLPKRPTSLPLNTKNSTKEPRLKFGSKHKSNLKQVE<br>TGVAKMNTINAAEPHIVTVTMNGVAGRNHSVNSHTATTQYANGVVPSGQTANTVAHRAQEMLQNQF<br>IGEDTRLNINSSPDEHEPLLRREQQAGHDEGVLDRLVDRRERPLEGGRTNSNNNNSNPCSEQEVPT<br>QGVPSTVADPGPSKPRRAQRPNSLDLSATNVLDGSSLQLGDSTQDGKSGSGEKIKKRVKTPYSLKR<br>WRPSTWVISTEPLDCEVNNNGKDRAVHSKSSTTVYLADGGTATTMVSKDIGMNCL* |
| **#2 Allele 2 (-4,+3)** | MTSSPRRPRRVPSLLWTVLLVSAAAASQNQERLCAFKDPYQQDLGIGESRISHENGTILCSKGSTC<br>YGLWEKSKGDINLVKQGCWSHIGDPQECHYEECVVTTTPPTFRMERTGSAAVARTCVMSTLLRIFH<br>LQTQHHSVHLIHLTEMRQ* |
| **#3 Allele 3 (+1)** | MTSSPRRPRRVPSLLWTVLLVSAAAASQNQERLCAFKDPYQQDLGIGESRISHENGTILCSKGSTC<br>YGLWEKSKGDINLVKQGCWSHIGDPQECHYEECVVTTTPPSNSEWNVPVLLL* |
| **#4 Allele 2 (-49)** | MTSSPRRPRRVPSLLWTVLLVSAAAASQNQERLCAFKDPYQQDLGIGESRISHENGTILCSKGSTC<br>YGLWEKSKGDINLVKQGCWSHIGDPQECHYEECVVTTTPPSMSTLLRIFHLQTQHHSVHLIHLTEM<br>RQ* |
| **#4 Allele 3 (-7) and #6 Allele 2 (-7)** | MTSSPRRPRRVPSLLWTVLLVSAAAASQNQERLCAFKDPYQQDLGIGESRISHENGTILCSKGSTC<br>YGLWEKSKGDINLVKQGCWSHIGDPQECHYEECVVTTTPPSMERTGSAAVARTCVMSTLLRIFHLQ<br>TQHHSVHLIHLTEMRQ* |
| **#5 Allele 2 (-9,+1)** | MTSSPRRPRRVPSLLWTVLLVSAAAASQNQERLCAFKDPYQQDLGIGESRISHENGTILCSKGSTC<br>YGLWEKSKGDINLVKQGCWSHIGDPQECHYEECVVTTTPPSTNVPVLLL* |
| **#5 Allele 2 (-15)** | MTSSPRRPRRVPSLLWTVLLVSAAAASQNQERLCAFKDPYQQDLGIGESRISHENGTILCSKGSTC<br>YGLWEKSKGDINLVKQGCWSHIGDPQECHYEECVVT/IQNGTYRFCCCSTDLCNVNFTENFPPPDT<br>TPLSPPHSFNRDETIIIALASVSVLAVLIVALCFGYRMLTGDRKQGLHSMNMMEAAASEPSLDLDN<br>LKLLELIGRGRYGAVYKGSLDERPVAVKVFSXANRQNFINEKNIYRVPLMEHDNIARFIVGDERVT<br>ADGRMEYLLVMEYYPNGSLCKYLSLHTSDWVSSCRLAHSVTRGLAYLHTELPRGDHYKPAISHRDL |

| | |
|---|---|
| | NSRNVLVKNDGTCVISDFGLSMKLTGNXLVRPGEEDNAAISEVGTIRYMAPEVLEGAVNLRDCESA<br>LKQVDMYALGLIYWEIFMRCTDLFPGXSVPEYQMAFQTEVGNHPTFEDMQVLVSREKQRPKFPEAW<br>KENSLAVRSLKETIEDCWDXDAEARLTAQCAEERMAELMMIWERNKSVSPTVNPMSTAMQNERNLS<br>HNRRVPKIGPYPDYSSSSYIEDSIHHTDSIVKNISSEHSMSSTPLTIGEKNRNSINYERQQAQARI<br>PSPETSVTSLSTNTTTTNTTGLTPSTGMTTISEVPYPDETSLHATNVSQPVGPTPVCLQLTEEDLE<br>TNKLDPKEVDKNLKESSDENLMEHSLKQFSGPDPLSSTSSSLPYPLIKLAVEVTGQQDFTQAANGQ<br>ACLIPDVPPTQIYPLPKQQNLPKRPTSLPLNTKNSTKEPRLKFGSKHKSNLKQVETGVAKMNTINA<br>AEPHIVTVTMNGVAGRNHSVNSHTATTQYANGVVPSGQTANTVAHRAQEMLQNQFIGEDTRLNINS<br>SPDEHEPLLRREQQAGHDEGVLDRLVDRRERPLEGGRTNSNNNNSNPCSEQEVPTQGVPSTVADPG<br>PSKPRRAQRPNSLDLSATNVLDGSSLQLGDSTQDGKSGSGEKIKKRVKTPYSLKRWRPSTWVISTE<br>PLDCEVNNNGKDRAVHSKSSTTVYLADGGTATTMVSKDIGMNCL* |
| **#8 Allele 2 (+94)** | MTSSPRRPRRVPSLLWTVLLVSAAAASQNQERLCAFKDPYQQDLGIGESRISHENGTILCSKGSTC<br>YGLWEKSKGDINLVKQGCWSHIGDPQECHYEECVVTTTPPSIQNGTYRFCCCSTDLCNVNYEEMCS<br>NYHPTINSEWNVPVLLL* |
